# Genome-engineering with CRISPR-Cas9 in the mosquito *Aedes aegypti*

**DOI:** 10.1101/013276

**Authors:** Kathryn E. Kistler, Leslie B. Vosshall, Benjamin J. Matthews

## Abstract

The mosquito *Aedes aegypti* is a potent vector of the Chikungunya, yellow fever, and Dengue viruses, which result in hundreds of millions of infections and over 50,000 human deaths per year. Loss-of-function mutagenesis in *Ae. aegypti* has been established with TALENs, ZFNs, and homing endonucleases, which require the engineering of DNA-binding protein domains to generate target specificity for a particular stretch of genomic DNA. Here, we describe the first use of the CRISPR-Cas9 system to generate targeted, site-specific mutations in *Ae. aegypti*. CRISPR-Cas9 relies on RNA-DNA base-pairing to generate targeting specificity, resulting in cheaper, faster, and more flexible genome-editing reagents. We investigate the efficiency of reagent concentrations and compositions, demonstrate the ability of CRISPR-Cas9 to generate several different types of mutations via disparate repair mechanisms, and show that stable germ-line mutations can be readily generated at the vast majority of genomic loci tested. This work offers a detailed exploration into the optimal use of CRISPR-Cas9 in *Ae. aegypti* that should be applicable to non-model organisms previously out of reach of genetic modification.

## Introduction

As a primary vector of the serious and sometimes fatal Chikungunya, yellow and Dengue viruses, the mosquito *Aedes aegypti (Ae. aegypti)* is responsible for hundreds of millions of human infections annually [1]. To transmit disease, a female mosquito must first bite an infected individual, and, after a period of viral incubation within the mosquito, bite another human. Female mosquitoes use cues such as odor, carbon dioxide, and temperature to locate a host and obtain a blood-meal [2], used to produce a clutch of approximately 100 eggs. Once a mosquito has developed mature eggs, she uses volatile and contact cues to locate and evaluate a body of water at which to lay her eggs, or oviposit. Our long-term goals involve using genome-engineering techniques coupled with quantitative behavioral analysis to investigate the genetic and neural basis of innate chemosensory behaviors in this important disease vector.

Clustered regularly interspaced palindromic repeats (CRISPR) and CRISPR associated (Cas) genes are components of an adaptive immune system that are found in a wide variety of bacteria and archaea [3]. Beginning in late 2012 [4], the bacterial type II CRISPR-Cas9 system has been adapted as a genome-engineering tool in a wide variety of organisms and *in vitro* preparations, dramatically expanding the ability to introduce specific genome modifications [3]. In particular, the ease of designing and generating these reagents at the bench has opened the door for studies of gene function in non-traditional model organisms.

The genome of *Ae. aegypti* is relatively large and incompletely mapped [5–8], presenting difficulties in recovering mutations generated by traditional forward genetics. *Ae. aegypti* has a recent history of genetic modification, including transposon-mediated transgenesis [9, 10] and loss-of-function gene editing with zinc-finger nucleases (ZFNs) [2, 11, 12], TAL-effector nucleases (TALENS) [13, 14], and homing endonuclease genes (HEGs) [15]. ZFNs and TALENs are modular DNA-binding proteins tethered to a non-specific FokI DNA nuclease [16], while HEGs are naturally occurring endonucleases that can be reengineered to target novel sequences [17]. Targeting specificity in these classes of genome-engineering reagents is conferred by context-sensitive protein-DNA binding interactions that are not completely understood. It is therefore technically difficult to engineer these proteins to target many sequences of interest.

In this paper, we describe methods for site-directed mutagenesis in *Ae. aegypti* using RNA-guided endonucleases (RGENs) based on the type II CRISPR-Cas9 system. RNA-DNA Watson-Crick base pairing is used to target the double-stranded endonuclease Cas9, derived from *Streptococcus pyogenes*, to specific genome locations where it introduces a double-stranded break. In just two years, the CRISPR-Cas9 system has been adapted for precision genome-engineering in dozens of model organisms from bacteria to primates [3, 18]. While many of these efforts have influenced our work, two studies in the vinegar fly *Drosophila melanogaster* [19] and the zebrafish *Danio rerio* [20] were particularly important in guiding our early attempts to adapt CRISPR-Cas9 to the mosquito *Ae. aegypti*.

A detailed bench manual with step-by-step guidance for designing, generating, and testing these reagents is available as a supplement to this paper. This work is not an exhaustive exploration of the parameters controlling the efficiency of CRISPR-Cas9 mutagenesis in *Ae. aegypti*, but is instead a practical guide for generating mosquito mutants. Given the proven flexibility of the CRISPR-Cas9 system, we believe that the protocols and procedures outlined here and by numerous other laboratories will continue to be optimized and modified for use in many organisms for which precision genome-engineering has not yet been employed.

## Materials and Methods

A detailed bench manual is available as Supplemental File S1.

### Microinjection and insect rearing

Microinjection into *Ae. aegypti* embryos was performed according to standard protocols [21] at the Insect Transformation Facility at the University of Maryland. Embryos were hatched 3 days post-injection and reared to pupal or adult stages according to previously described rearing procedures [2,12]. Liverpool IB12 (LVP-IB12) mosquitoes [5] were used both as the injection strain and as the wild-type strain for out-crossing and were obtained through the MR4 as part of the BEI Resources Repository, NIAID, NIH. This *Ae. aegypti* LVP-IB12, MRA-735 strain, was originally deposited by M.Q. Benedict. Female mosquitoes were provided with a mouse or human blood source for egg production. All laboratory blood-feeding procedures with mice and humans were approved and monitored by The Rockefeller University Institutional Animal Care and Use Committee and Institutional Review Board, protocols 11487 and LVO-0652, respectively. All humans gave their informed consent to participate in mosquito blood-feeding procedures.

### Cas9 mRNA and protein

Cas9 mRNA was transcribed from vector pMLM3613 (plasmid 42251, AddGene) [20] using mMessage mMachine T7 Ultra Transcription kit (AM1345, Life Technologies). Recombinant Cas9 protein was obtained commercially (CP01, PNA Bio).

### sgRNA design and construction

sgRNAs were designed by manually searching genomic regions for the presence of protospacer-adjacent motifs (PAMs) with the sequence NGG, where N is any nucleotide. We required that sgRNA sequences be 17-20 bp in length, excluding the PAM, and contain one or two 5’ terminal guanines to facilitate transcription by T7 RNA polymerase. sgRNA sequences were checked for potential off-target binding using the following two web tools: http://zifit.partners.org/ZiFiT/ and http://crispr.mit.edu. To minimize the potential for off-target mutagenesis, we avoided sgRNA sequences with closely related binding sites, defined as fewer than 3 mismatched bases to reduce the possibility of cutting, whenever possible. See Table 1 for sgRNA sequences and closest matches.

**Table 1.**
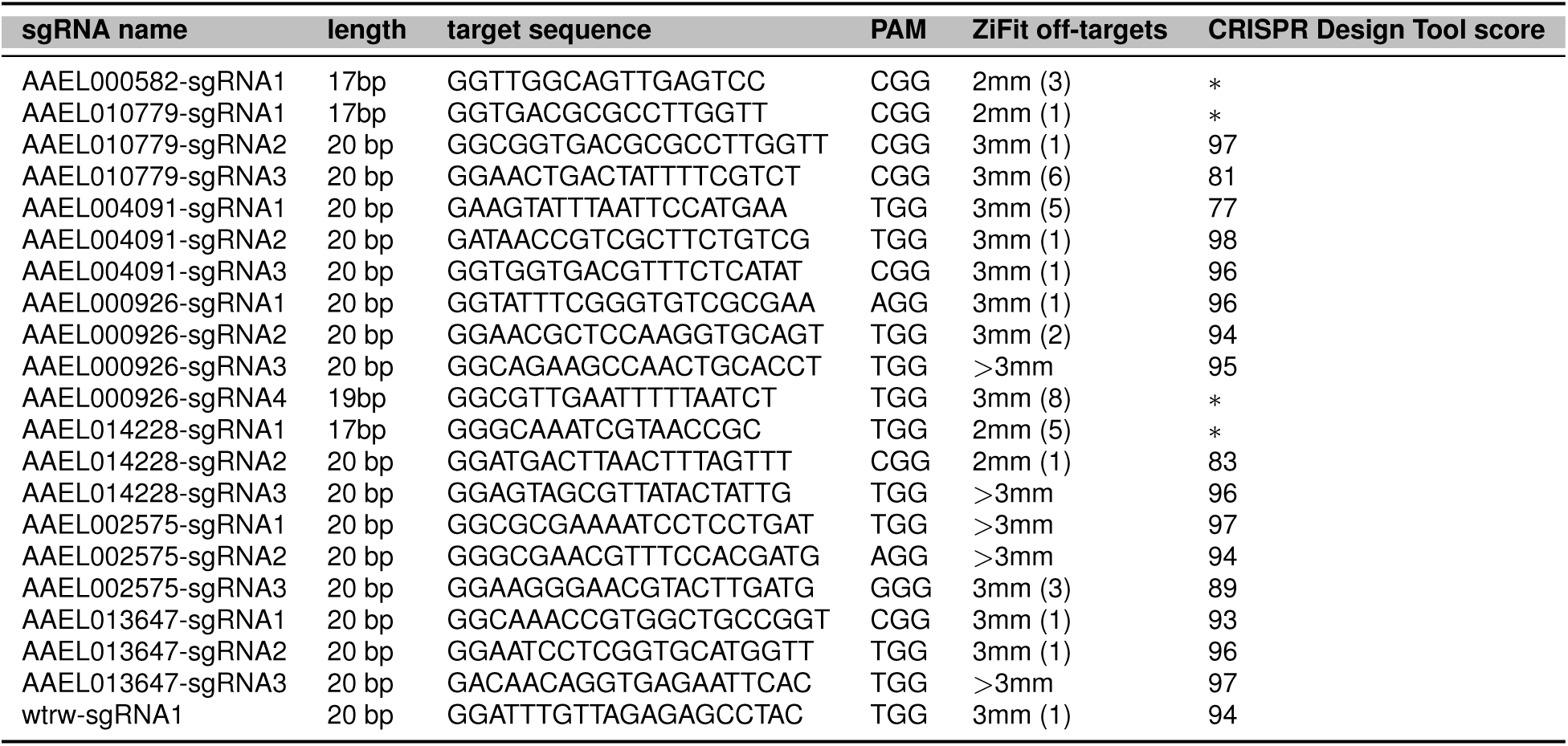
Sequences of sgRNAs used in this study. Potential off-target sites were evaluated by two publicly available tools. ZiFiT (http://zifit.partners.org) was used to determine the number of potential off-target sites in the *Ae. aegypti* genome with 3 or fewer mismatched bases (mm). The number of distinct sites with the given number of mismatches is noted in parentheses. The CRISPR Design Tool (http://crispr.mit.edu) uses an algorithm to evaluate potential off-target sites in the context of the *Ae. aegypti* genome and returns a specificity score for sgRNAs of length 20 bp. Scores above 50 are considered”high quality”. *indicates sgRNA targets of 17-19bp that cannot be tested with the CRISPR Design Tool

Linear double-stranded DNA templates for specific sgRNAs were produced by a template-free polymerase chain reaction (PCR) with two partially overlapping oligos (primers sgRNA-R and sgRNA-F; Table 2) [19]. In early experiments, sgRNA sequences were cloned into pDR274 [20] and linearized with PmeI (R0560, NEB). In both cases sgRNAs were produced using the MegaScript T7 *in vitro* transcription kit (AM1334, Life Technologies) overnight at 37°C. Transcribed sgRNA was purified with MegaClear columns (AM1908, Life Technologies).

**Table 2.**
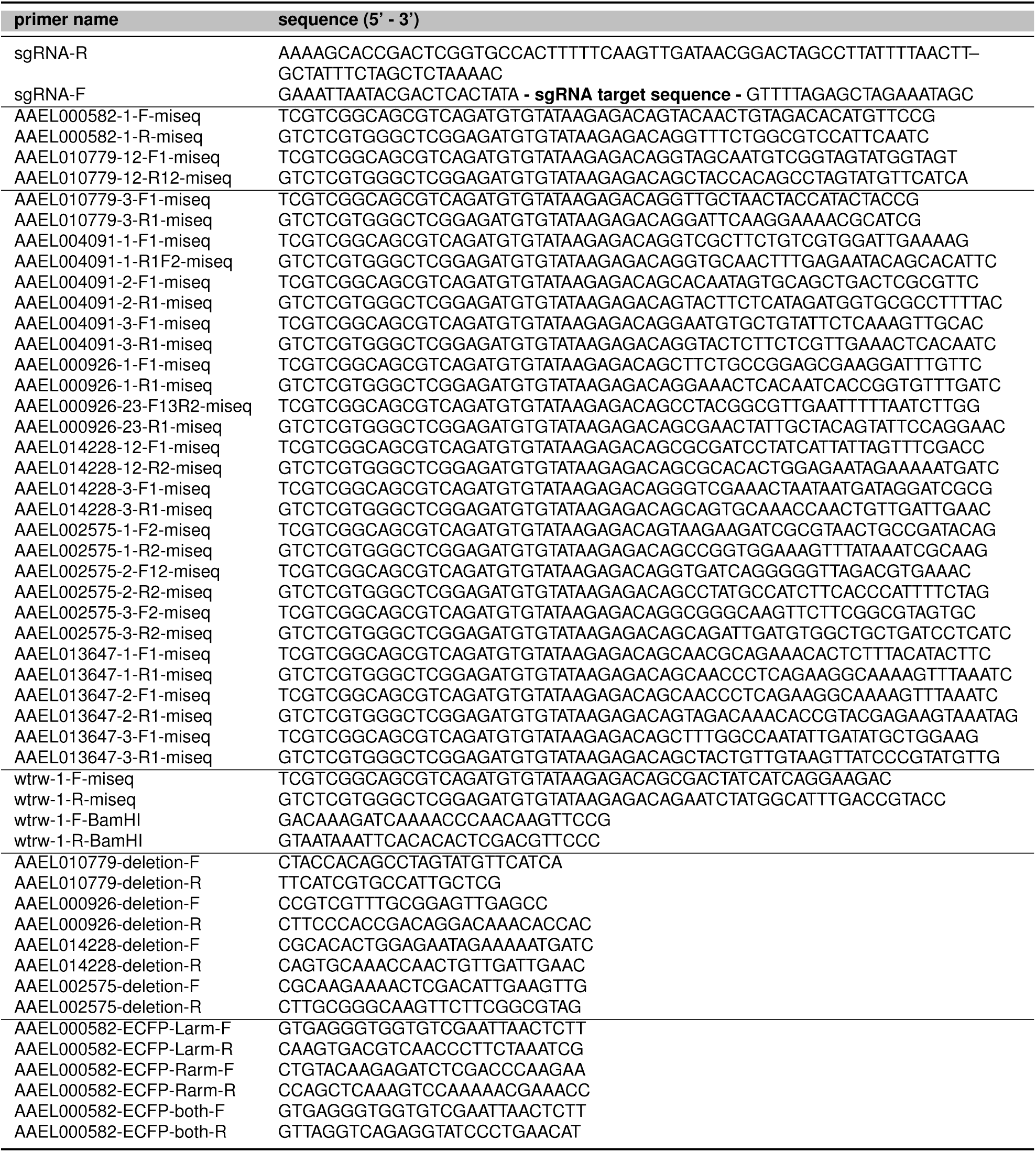
PCR primer sequences used in this study

### Extraction of genomic DNA

Genomic DNA was extracted from individual or pools of mosquitoes using either the DNEasy Blood and Tissue Kit (69581, Qiagen) or a 96-well plate extraction protocol [22].

### Sequencing and analysis of CRISPR-Cas9 induced mutations

A two-step PCR protocol was used to amplify amplicons surrounding the putative CRISPR-Cas9 cut site from genomic DNA of G_0_ mosquitoes (those that were injected as embryos and allowed to develop to pupal or adult stages) or G_1_ individuals (the progeny of G_0_ individuals crossed to LVP-IB12). First, genome-specific primers were designed with the following tails:

- `Forward: 5′-TCGTCGGCAGCGTCAGATGTGTATAAGAGACAG-GENESPECIFIC-FORWARD-3’`
- `Reverse: 5′-GTCTCGTGGGCTCGGAGATGTGTATAAGAGACAG-GENESPECIFIC-REVERSE-3’`

20 cycles of PCR was performed using KOD Hotstart polymerase (71086-3, EMD Millipore). The product of the first PCR was used as a template for a second round of PCR with Nextera XT Indexed Primers (item FC-131-1002, Illumina). PCR cycles were kept to a minimum to reduce the non-linearity of PCR amplification that can occur at higher cycle numbers. Samples were purified using Ampure XP magnetic beads at 0.6x volume to separate amplicons from primers, were examined on an Agilent Bioanalyzer to verify purity and sizing, and pooled in roughly equimolar amounts. To ensure high sequencing quality, final amplicon pools were quantitated by qPCR to determine the precise molarity of amplicons containing the adapters need for clustering and sequencing (KK4844, KAPA BioSystems). Samples were sequenced on an Illumina MiSeq following the manufacturer’s instructions.

Following sequencing, reads were aligned to reference sequences of the PCR amplicons and examined for the presence of insertions, deletions, or other polymorphisms. Scripts developed for the analysis of this data, including those used to run the alignments and mutation quantitation for all figures, are available at https://github.com/bnmtthws/crispr_indel. Other software packages used include GMAP/GSNAP [23] http://research-pub.gene.com/gmap/, GATK [24] https://www.broadinstitute.org/gatk/, and pysamstats https://github.com/alimanfoo/pysamstats.

### Donor construction for homology-directed repair

Single-stranded DNA oligodeoxynucleotides (ssODNs) were ordered as 200 bp ‘Ul-tramers’ from Integrated DNA Technologies (IDT). Plasmids used as double-stranded DNA donors were constructed by PCR-amplifying homology arms from LVP-IB12 genomic DNA and cloning with In-Fusion HD cloning (Clontech) into one of two base plasmids, PSL1180polyUBdsRED (AddGene #49327) and pSL1180-HR-PUbECFP (AddGene #47917). Annotated sequences of oligonucleotides and plasmids used for homology-directed repair are available as Supplemental File S2.

### Molecular genotyping of stable germ line alleles by PCR

To verify the presence of exogenous sequences inserted by homology-directed-repair or the presence of insertions and deletions, PCR amplicons surrounding the putative cut site were generated from genomic DNA (see Table 2 for primer sequences). Following purification, amplicons were Sanger sequenced (Genewiz), or used as a template for a restriction digest using the enzyme BamHI (R0136, New England Biolabs [NEB]) or PacI (R0547, NEB).

### Genotyping stable germ line alleles by fluorescence

Larvae or pupae were immobilized on a piece of moist filter paper and examined under a dissection microscope (SMZ1500, Nikon) with a fluorescent light source and ECFP and dsRed filtersets.

## Results

### Outline of CRISPR and injection components

The bacterial type-II CRISPR system has been adapted in many organisms to generate RNA-guided endonucleases, or RGENs, that are targeted to specific regions of the genome by RNA-DNA base-pairing [3]. The core of this adapted CRISPR/Cas system is comprised of two components: 1) a synthetic single guide RNA (sgRNA), which is a small RNA containing 17-20 bases of complementarity to a specific genomic sequence, and 2) the Cas9 nuclease derived from *Streptococcus pyogenes* (SpCas9). SpCas9 forms a complex with the sgRNA and induces double-stranded DNA breaks at regions of the genome that fulfill two criteria: 1) contain a sequence complementary to the recognition site of the sgRNA that is 2) directly adjacent to a protospacer-adjacent motif (PAM). The PAM sequence for SpCas9 takes the form of NGG. These motifs occur approximately once every 17 bp in the *Ae. aegypti* genome, making it possible to target essentially any genomic loci of interest with the CRISPR-Cas9 system.

To generate stable mutations that can be transmitted through the germline, CRISPR-Cas9 reagents must be introduced pre-blastoderm stage embryos composed of a syncytium of nuclei prior to cellularization (Figure 1). This early developmental stage offers access of genome-editing reagents to the nuclei of both somatic and germ-line cells. In brief, embryos are microinjected 4-8 hours after egg-laying, and allowed to develop for 3 days before being hatched in a deoxygenated hatching solution [21]. Following hatching, pupae are collected for sequencing or allowed to emerge as adults and outcrossed to wild-type LVP-IB12 males or females, as appropriate. Genomic DNA from injected embryos is examined for modification rates. Following blood-feeding, G_1_ eggs are collected from these outcrosses to screen for germ-line transmission of stable mutations.

**Figure 1.**
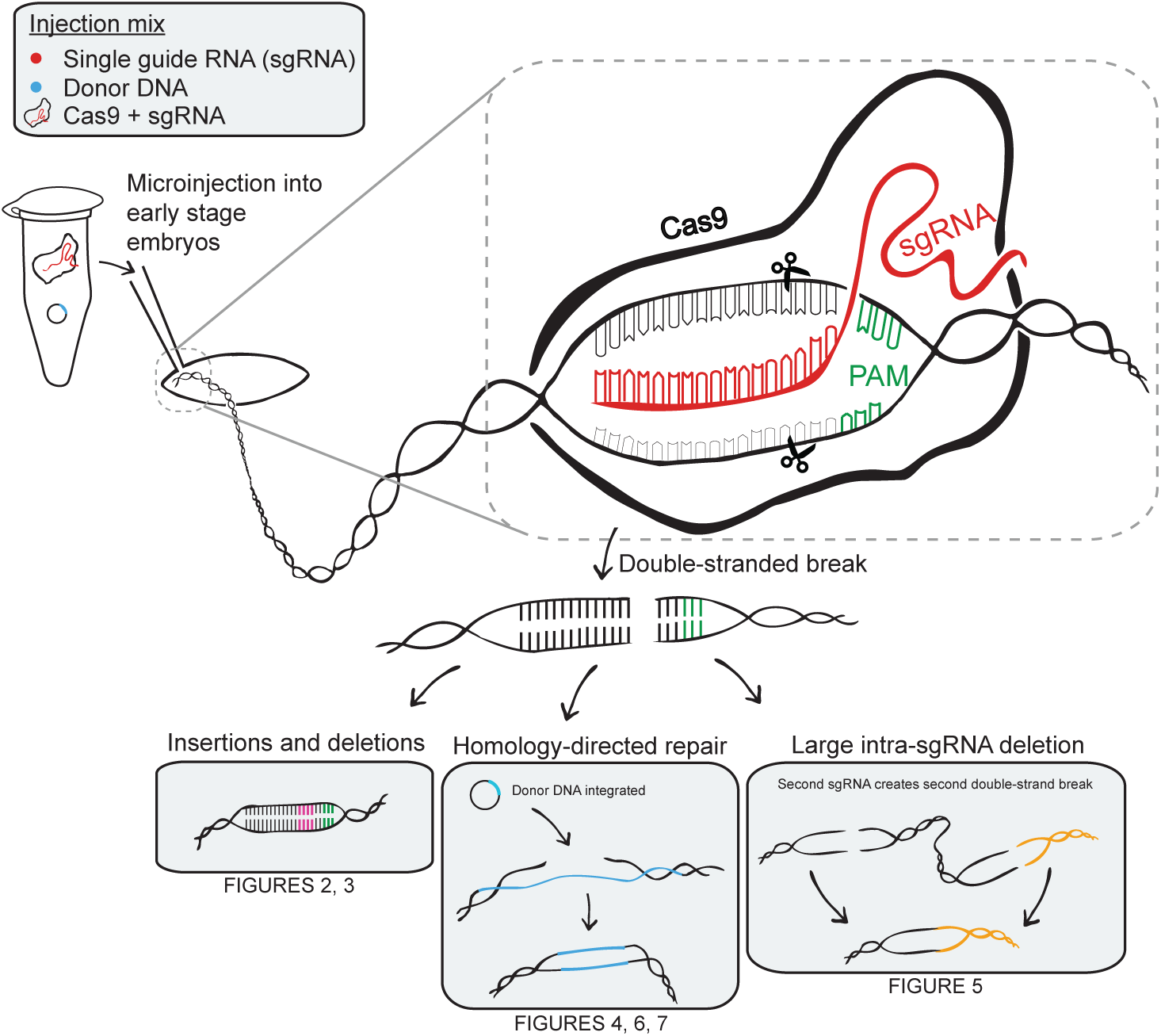
**CRISPR-Cas9 genome-engineering system as adapted to the mosquito *Ae. aegypti***. An sgRNA, donor DNA, and the nuclease Cas9 are microinjected into *Ae. aegypti* embryos. A 17-20 bp sequence of the sgRNA binds to the complementary site in the genome, targeting Cas9 and generating a double-stranded break. These breaks can be repaired in a number of ways, including the following: insertions or deletions resulting from error-prone non-homologous end-joining, insertion of exogenous DNA sequences through homology-directed repair, and generating large deletions between multiple sgRNA/Cas9 double-stranded breaks.

When faced with a double-stranded DNA break, DNA repair machinery can resolve this break in one of two ways: non-homologous end-joining, which can result in small insertions and deletions, or, less frequently, homology-directed repair, which uses exogenous sequence containing regions of homology surrounding the cut site as a template for repair. Additionally, cutting with multiple sgRNAs can result in large deletions between the two cut sites. In this paper, we will discuss stable germ-line transmission of all three types of these alleles in *Ae. aegypti*.

### Identifying optimal injection mixes for CRISPR-Cas9 mutagenesis

The insertions and deletions resulting from non-homologous end-joining act as a signature for evaluating the efficiency of reagents such as ZFNs, TALENs, and CRISPR-Cas9. Quantifying the level of these insertions and deletions can give an indication of the activity of a particular sgRNA/Cas9 combination. A variety of techniques have been described to evaluate the cutting efficiency of Cas9/sgRNA RNA-guided nucleases. These include enzymatic detectors of polymorphisms such as Surveyor or T7 Endonuclease I [25], high-resolution melting point analysis (HRMA) [26], Sanger sequencing [27] or deep sequencing [28]. Each of these techniques evaluates the level of polymorphism in a short PCR-generated amplicon surrounding the sgRNA target site. With the exception of deep-sequencing, these approaches provide only semi-quantitative estimates of the effectiveness of mutagenesis in each sample. In addition, the *Ae. aegypti* genome is highly polymorphic, and so techniques that are based on detecting sequence mismatches, such as HRMA, T7E1, or Surveyor nuclease assays, are prone to false positives from polymorphisms present in our wild-type strains.

For these reasons, we opted to prepare and deep sequence barcoded PCR amplicons surrounding putative CRISPR-Cas9 cut sites from small pools of injected animals to accurately determine the rates of cutting at different genomic loci. Briefly, sequencing libraries are prepared using a two-step PCR process that incorporates adapter and barcode sequences necessary for Illumina sequencing (Figure 2A). Many distinct barcoded amplicons can be pooled together using Illumina-supplied Nextera XT primers. We estimate that 10,000-100,000 reads are ample for this analysis, meaning that at current prices, sequencing of amplicons from 3 sgRNAs per gene, for 10 distinct genes can be run for a cost of approximately $70 per gene (MiSeq v3 reagents, 150-cycle flowcell, item MS-102-3001).

**Figure 2.**
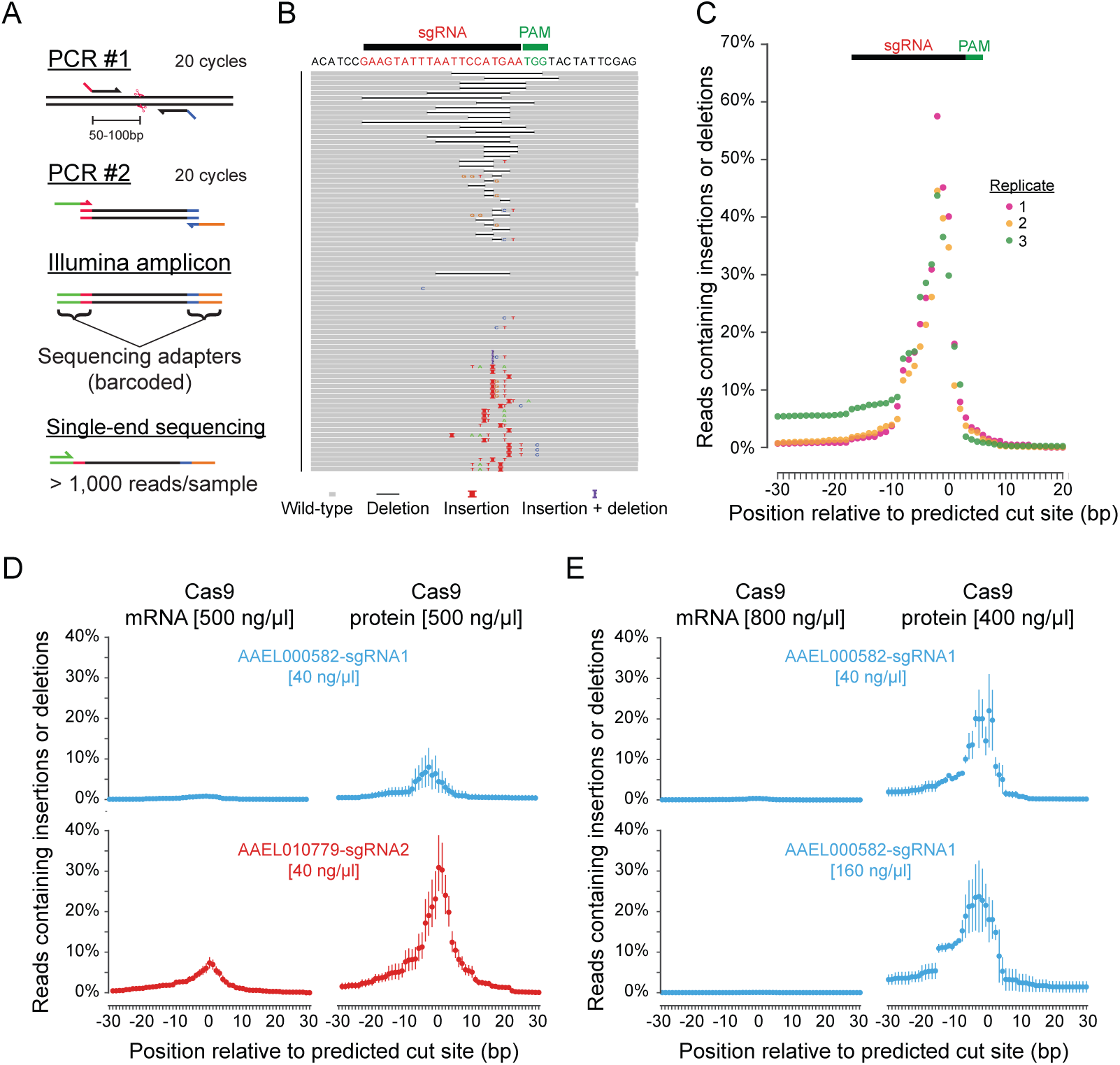
**Deep sequencing approach to quantifying CRISPR-Cas9 efficiency A)** Schematic of strategy to generate barcoded PCR amplicons for sequencing. In this example (B-C) amplicons were generated from adults reared from embryos injected with Cas9 and an sgRNA (AAEL004091-sgRNA1) using primers AAEL004091-1-F and AAEL004091-1-R. **B)** Visualization of a subset of alignments to the amplicon reference sequence (top) reveals a heterogeneity of insertions, deletions, and other changes. **C)** Quantification of three replicate libraries, plotted as a percentage of reads aligned to a given base that contain an insertion or deletion at that base, shows reproducibility of mutation rates between individual pools of pupae derived from the same injection. Data are represented as a percentage of reads aligned to a given base that contain an insertion or deletion at that base, and is plotted as mean (circle) and 95% confidence intervals (line). **D)** Summary of sequencing data from animals injected with two different sgRNAs in combination with Cas9 mRNA (left) or Cas9 recombinant protein (right), both at 500 ng/μL. **E)** Increasing Cas9 mRNA concentration to 800 ng/μL (left) did not improve cut rate in conjunction with an sgRNA at 40 or 160 ng/μL. Reducing Cas9 protein concentration to 400 ng/μL (right) increased insertion or deletion rate slightly when combined with an sgRNA at 40 ng/μL. Increasing sgRNA concentration to 160 ng/μL did not dramatically affect the insertion or deletion rate.

Following sequencing, reads were aligned to a reference sequence using the GSNAP short read aligner [23], and insertions and deletions were quantified using the python package pysamstats. This procedure results in data on the number of polymorphisms, including insertions and deletions, found in reads that span each nucleotide of a reference sequence (Figure 2B). Notably, in injections containing a sgRNA and Cas9, a pattern of elevated insertions and deletions can be observed with a peak 3 bp 5’ of the beginning of the PAM, exactly the position at which Cas9 is known to make a double-stranded break (Figure 2C). Importantly, there was high concordance in the mutagenesis rates seen between multiple biological replicates generated from different pools of pupae from a single injection (Figure 2C), suggesting that there is minimal variability in mutagenesis frequency in individual injected animals.

We next varied the delivery method and concentration of Cas9 and the concentration of a given sgRNA to determine an optimal injection mix composition that would yield high levels of somatic and germline mutations. We first attempted mutagenesis with plasmid DNA constructed to express Cas9 under the control of an *Ae. aegypti* polyubiquitin (PUb) promoter that can effectively drive TALEN half-sites when microinjected into embryos [13,29]. However, injection mixes containing PUb-Cas9 plasmid and a validated sgRNA did not induce detectable rates of insertions or deletions (data not shown).

The failure of Cas9 expressed by plasmid DNA might be due to the temporal dynamics of transcription and translation, so we explored two additional methods of Cas9 delivery: mRNA produced by *in vitro* transcription and recombinant SpCas9 protein. When included at 500 ng/μL, both Cas9 mRNA and protein induced detectable mutagenesis at two distinct guide RNA sites. However, recombinant Cas9 protein induced mutagenesis at rates 5-10x higher than Cas9 mRNA (Figure 2D). To test whether the concentration of sgRNA or Cas9 mRNA or protein was limiting in these earlier injections, we tried four additional injection mixes (Figure 2E), with a single validated sgRNA, and determined that mixes containing 400 ng/μL Cas9 recombinant protein induced the highest rates of mutagenesis. Increasing sgRNA concentration did not dramatically increase mutagenesis rates.

### Identifying active sgRNAs for a given genomic target

We reasoned that sgRNAs that show higher somatic cut rates in injected animals would be more likely to result in stable germ-line transformation. We first designed 3 different sgRNAs against 6 different genes using publicly-accessible design tools to minimize the potential for off-target mutagenesis (Figure 3A). These varied in length between 17-20 bp [30], and in addition to being adjacent to a PAM sequence, were required to begin with GG or G to facilitate *in vitro* transcription of sgRNA.

**Figure 3.**
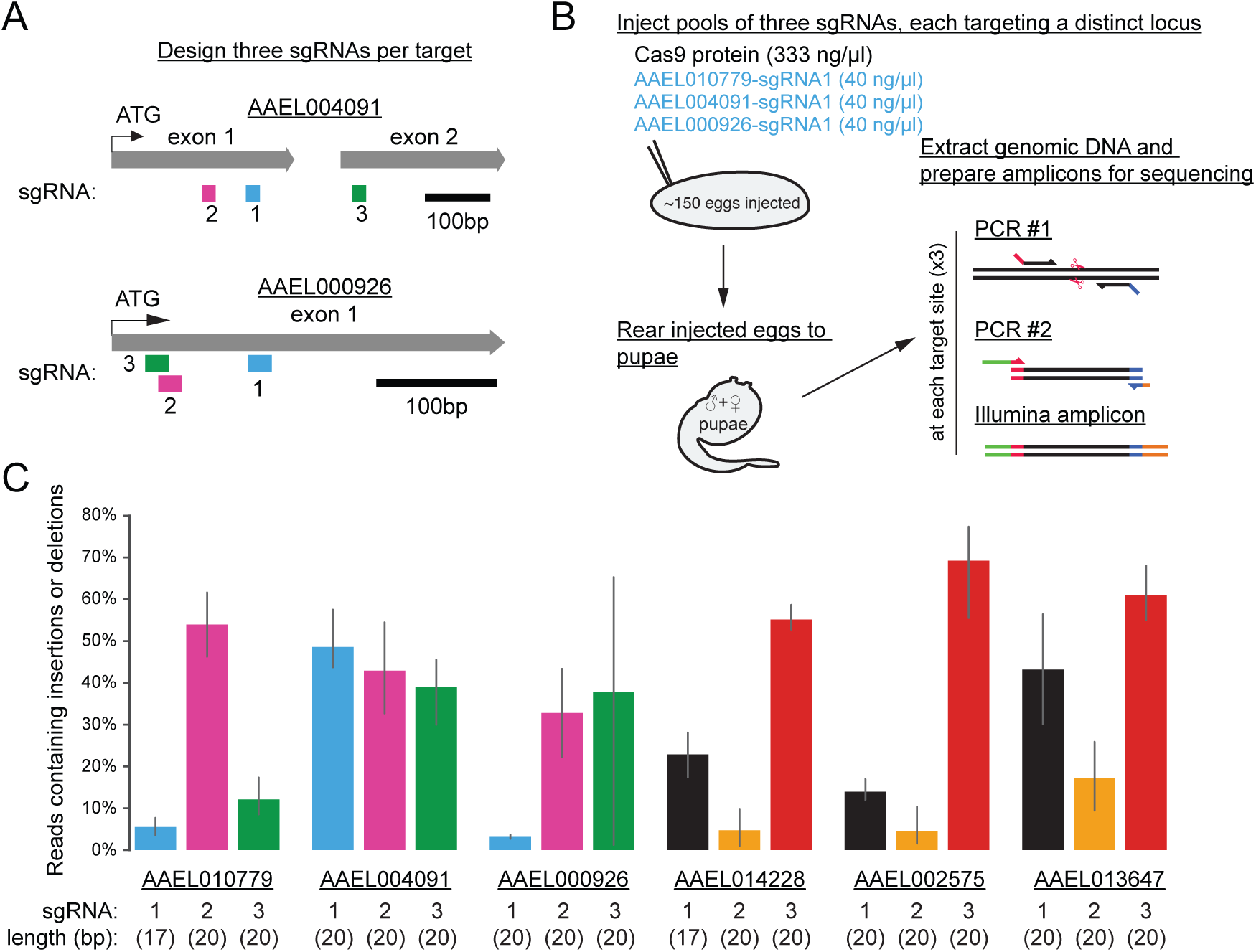
**Identifying active sgRNAs A)** Schematic of two target genes in the *Ae. aegypti* genome: AAEL004091 and AAEL000926. Three sgRNAs were designed against the first 500 bp of each gene. **B)** Schematic of the workflow for a small injection (approx. 150 embryos) of Cas9 protein and a pool of three sgRNAs against three distinct target genes. After rearing to pupal stages to synchronize developmental time, genomic DNA was extracted and sequencing amplicons were prepared against each of the three targets. **C)** Sequencing results from 6 small injections (sgRNA sequences can be found in Table 1).

To test the efficiency of these sgRNAs, we performed a series of 6 small test injections (145-168 embryos each) into *Ae. aegypti* embryos (Figure 3B). Each injection mix was comprised of recombinant Cas9 protein at 333 ng/μL and a pool of three sgRNAs (40 ng/μL each), each targeted against a different gene. Survival rates were very high for these injections (ranging from 46.1%-63.3%, as compared to a 18.6% average survival in previous experiments with 500 ng/μL Cas9 protein or mRNA), and we attribute this marked increase in survival to the reduction in Cas9 (or overall injection mix) concentration. Surviving embryos were reared to pupal stages and collected for PCR amplicon preparation and sequencing analysis (Figure 3B).

All 18 sgRNAs induced detectable levels of mutagenesis, although the rates of cutting were highly variable within and between different genomic targets (Figure 3C). Thus, in addition to potential effects of chromatin state or other genome accessibility at a particular genomic locus, there appear to be sequence- or context-dependent effects on sgRNA efficiency that are not yet fully understood. Designing and testing 3 sgRNAs resulted in the identification of at least one highly active sgRNA for the 6 genomic targets tested here. The variability observed between different sgRNAs targeting the same locus underscores the benefits of testing multiple sgRNAs per gene before undertaking large-scale mutagenesis injections.

### Germ-line transmission of mutant alleles

We next examined whether somatic mutagenesis detected in adults reared from injected embryos (G_0_ animals) could result in transmission of stable mutant alleles through the germline to their offspring (G_1_ animals). We designed an sgRNA near the 5’ end of *Aaeg-wtrw*, a single-exon gene identified through bioinformatics which bears homology to the *D. melanogaster* TRP channel *water witch (wtrw)* [31]. We also included 300 ng/μL of a 200 bp ssODN donor to act as a template for homology-directed repair. The ssODN comprised homology arms of 87-90 bases on either side of the Cas9 cut site, with an insert containing stop codons in all three frames of translation as well as an exogenous restriction enzyme recognition site. Successful integration of this template would result in a truncated protein of 91 amino acids rather than a full-length protein of 908 amino acids (Figure 4A).

**Figure 4.**
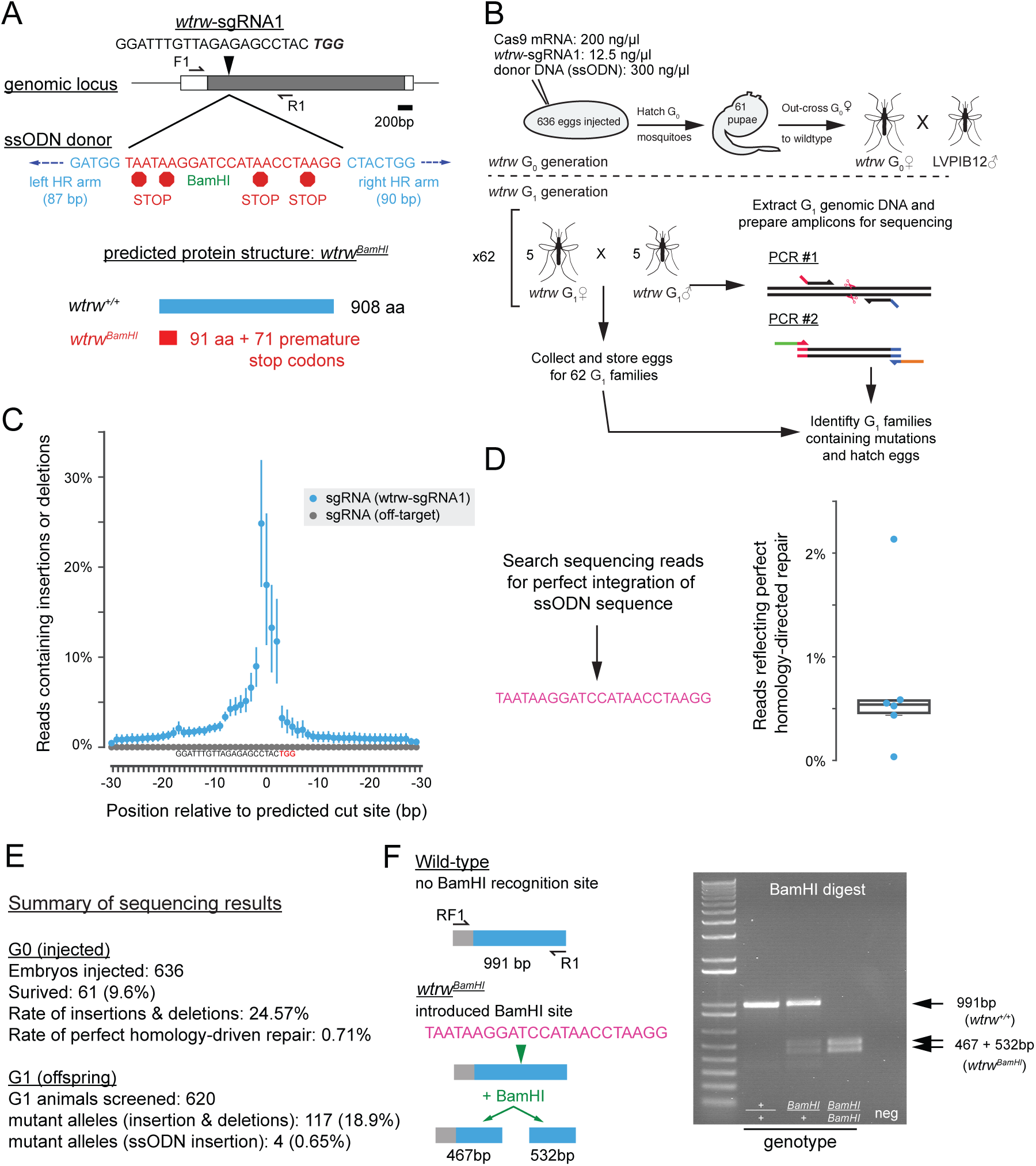
**Germ-line transmission of mutant alleles A)** Schematic of the *Ae. aegypti* wtrw locus detailing the sgRNA binding site wtrw-sgRNA1, the sequence of a 200 bp single-stranded oligonucleotide (ssODN) donor, and the predicted protein structure of a successfully modified locus. **B)** Schematic of injection performed to isolate mutations. Sequencing libraries were generated from both G_0_ and G_1_ animals after egg-laying. **C)** Summary sequencing data from G_0_ adults reveals robust levels of insertion and deletion in injections of wtrw-sgRNA1 and Cas9 mRNA. **D)** Exogenous ssODN sequence used to as a query for the unix tool grep to identify sequencing reads containing perfect homology-directed repair. A mean of 0.71% of reads contained this insertion. **E)** Summary statistics of sequencing reads from G_0_ and G_1_ individuals. **F)** Wild-type amplicons contain no BamHI recognition site while mutant alleles are cut into bands of 467 and 532 bp.

636 embryos were injected with a mixture of 200 ng/μL Cas9 mRNA and 12.5 ng/μL sgRNA. Note that these injections were performed prior to the optimization of injection mixes described above. To determine the activity of our injected reagents, we performed amplicon sequencing on 6 pools of 5-6 adult G_0_ animals after they were outcrossed and allowed to lay eggs (Figure 4B). After alignment and analysis of reads aligning to the reference sequence, we found that these samples contained a mean maximum somatic insertion or deletion rate of 24.87% centered on the Cas9 cut site (Figure 4C), indicating a high rate of somatic mosaicism at the targeted locus in animals injected with this cocktail. As a control, we sequenced amplicons at the *Aaeg-wtrw* locus from animals injected with an injection mix containing another sgRNA targeting a different region of the genome. These samples contained no detectable insertions or deletions at the *Aaeg-wtrw* locus (Figure 4C). Across the six samples, a mean of 0.71% of reads that aligned to the reference sequence contained sequences corresponding to the exogenous sequence in the single-stranded DNA donor (Figure 4D). This indicated that the ssODN template could drive homology-directed repair in somatic tissue, though at a much lower frequency than insertions or deletions mediated by non-homologous end-joining.

To determine whether these mutations were stably transmitted through the germline, we sequenced PCR amplicons derived from pools each containing 5 male and 5 female G_1_ offspring that were previously mated together and from which we had already collected F2 eggs. Analysis of resulting insertions and deletions using Genome Analysis Toolkit [GATK; [24]] revealed that there were at least 117 mutant chromosomes spread across 50 pools, meaning that the G_1_ mutation rate was at least 117/620, or 18.9% (Figure 4E). Four of these alleles corresponded perfectly to the sequence of the ssODN, meaning that our rate of stable germline transmission of alleles generated by homology-driven repair recombination is at least 0.6% (Figure 4E). Notably, these numbers correlate well with those derived from G_0_ animals (Figure 4E), suggesting that assays of somatic mutagenesis can be useful in predicting the efficiency of germ-line mutagenesis.

We hatched F2 eggs from a single family containing an allele generated by homology-directed repair. Sequencing of single-pair crosses allowed us to isolate a stable mutant line that can be genotyped by a simple restriction digest with the enzyme site introduced by the ssODN targeting event (Figure 4F). To enable future phenotypic characterization, this line was outcrossed for 8 generations to wild-type mosquitoes to increase genetic diversity in the mutant strain and to reduce the possibility of retaining off-target mutations in this population of animals.

### Deletions induced by multiplexed sgRNAs

Double-stranded breaks induced at multiple sgRNA sites can induce large deletions between the two cut sites in *D. melanogaster* [32]. We performed a series of five injections into small numbers of embryos using sgRNAs targeting 3 different genes. All but one of these sgRNAs (AAEL000926-sgRNA4) were previously validated as having high activity in somatic mutagenesis assays. We also included ssODN donors with arms on either side of the two breaks and an exogenous sequence containing stop codons in all 3 frames of translation as well as a restriction enzyme recognition site (Figure 5A). We injected small numbers of embryos and collected G_1_ embryos from all female G_0_ animals, which were hatched and screened individually (Figure 5B). From these individual families, we screened male pupae in pools via PCR, identifying potential mutations through size-based genotyping on an agarose gel. Individual females were outcrossed, blood-fed, and allowed to lay eggs. PCR amplicons were generated from these animals individually and examined for size-shifted bands on agarose gels (Figure 5C). Candidate alleles were characterized by Sanger sequencing of gel-purified fragments (Figure 5C). We found a wide range of mutant transmission rates in female G_1_ animals derived from single G_0_ individuals where genotyping male pupae showed the offspring of these G_0_ animals to contain at least one mutant G_1_ (Figure 5D).

**Figure 5.**
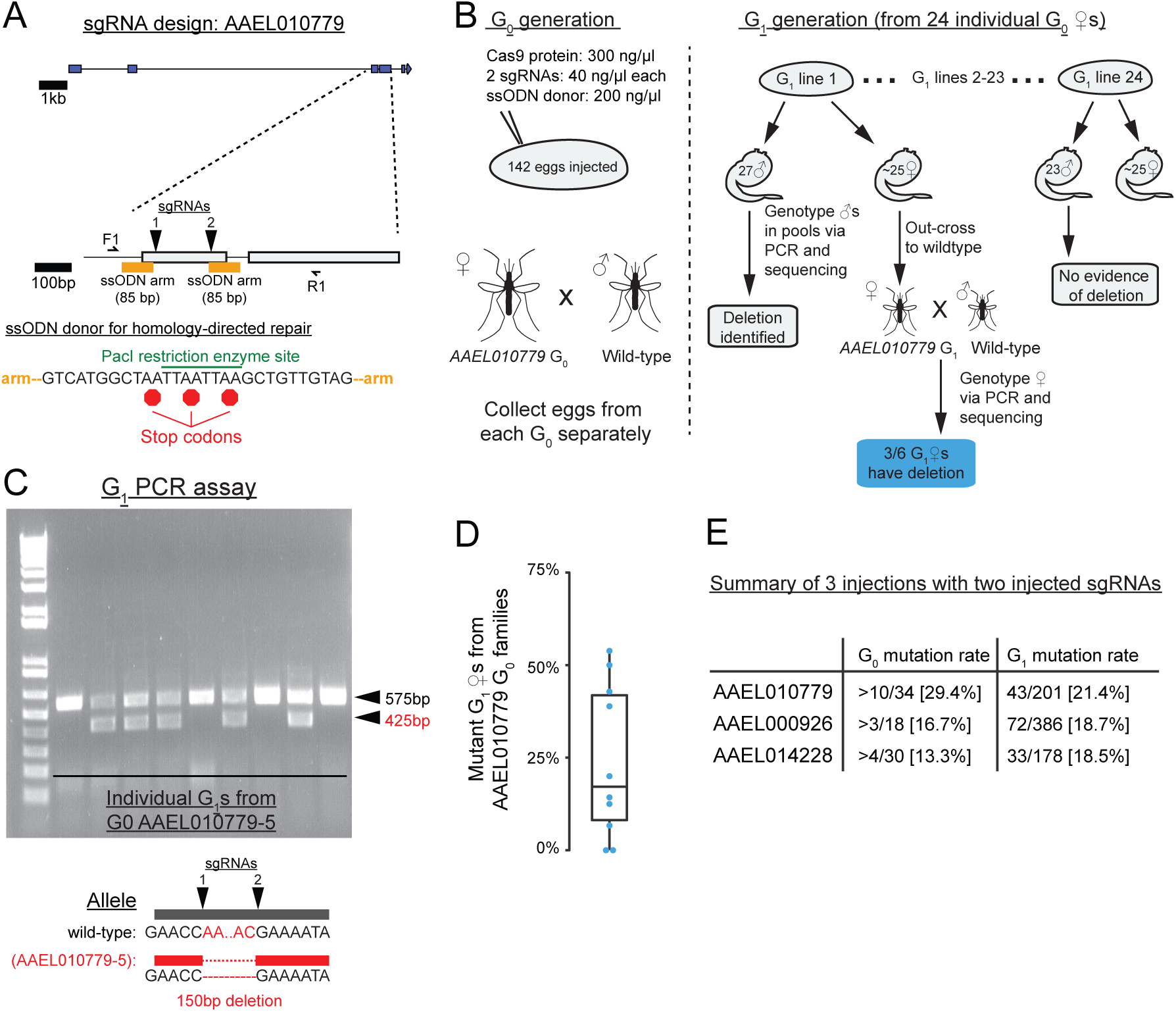
**Deletions generated by multiple sgRNAs A)** Schematic of the AAEL010779 genomic locus detailing the design of two sgRNAs and a ssODN donor. **B)** Injection strategy to identify deletion events in G_1_ animals with the injection of a small (125-150) number of embryos. **C)** Example agarose gel of 9 G_1_ females that are the offspring of a single G_0_ female. All individuals contain the wild-type band (black arrowhead). Sanger sequencing verified that the smaller band (red arrowhead) present in 5/9 heterozygous individuals is the result of a clean deletion between the two sgRNA cut sites. **D)** Plot of the proportion of mutant AAEL010779 G_1_ females from the 10 G_0_ families identified as containing at least one mutant allele. **E)** Summary data from 3 injections of this type show high rates of G_1_ mutagenesis across all 3 injections.

Although we did not fully characterize each potential mutant allele, we did note several different types of mutations in those that were characterized. These ranged from simple deletions, to homology-directed repair from the ssODN donor, to more complex modifications, including polymorphisms, inversions, and duplications. This indicates that induction of multiple double-stranded breaks is highly mutagenic. We also note that we were successful in obtaining germ-line mutations at high rates in all 3 injections (Figure 5E), making this a cost-effective and efficient way to generate loss-of-function mutant alleles. The relatively large size of deletions generated by this method makes sized-based molecular genotyping straightforward and greatly facilitates the out-crossing and homozygosing of mutant lines.

### Integration and transmission of large fluorescent cassettes

Finally, we asked whether CRISPR-Cas9 could be used to introduce longer cassettes of exogenous sequence via homology-dependent repair. In previous genome-engineering experiments in *Ae. aegypti*, zinc-finger nucleases were used to introduce large cassettes from a plasmid DNA donor with homology arms of at least 800 bp on either side [2, 11]. The insertion of a fluorescent cassette simultaneously creates a null mutant by interrupting the protein-coding sequence and inserts the visible fluorescent reporter. We performed injections with Cas9 protein, a validated sgRNA, and a plasmid donor containing homology arms of 799 bp and 1486 bp. This plasmid contains a cassette comprising the constitutive PUb promoter [29] driving the expression of a fluorescent reporter (Figures 6A-B). These arms were cloned from the strain of mosquito into which injections were performed to maximize homology, and were designed to avoid repetitive sequences such as transposable elements. Following injection, individual female G_0_ animals were outcrossed to wild-type mosquitoes and G_1_ eggs were collected (Figure 6A). Past experiences in our laboratory indicate that successful homology-directed repair occurs primarily, if not exclusively, in female G_0_ animals in *Ae. aegypti*. We therefore discarded G_0_ males and restricted our screening to G_1_ families generated from females.

**Figure 6.**
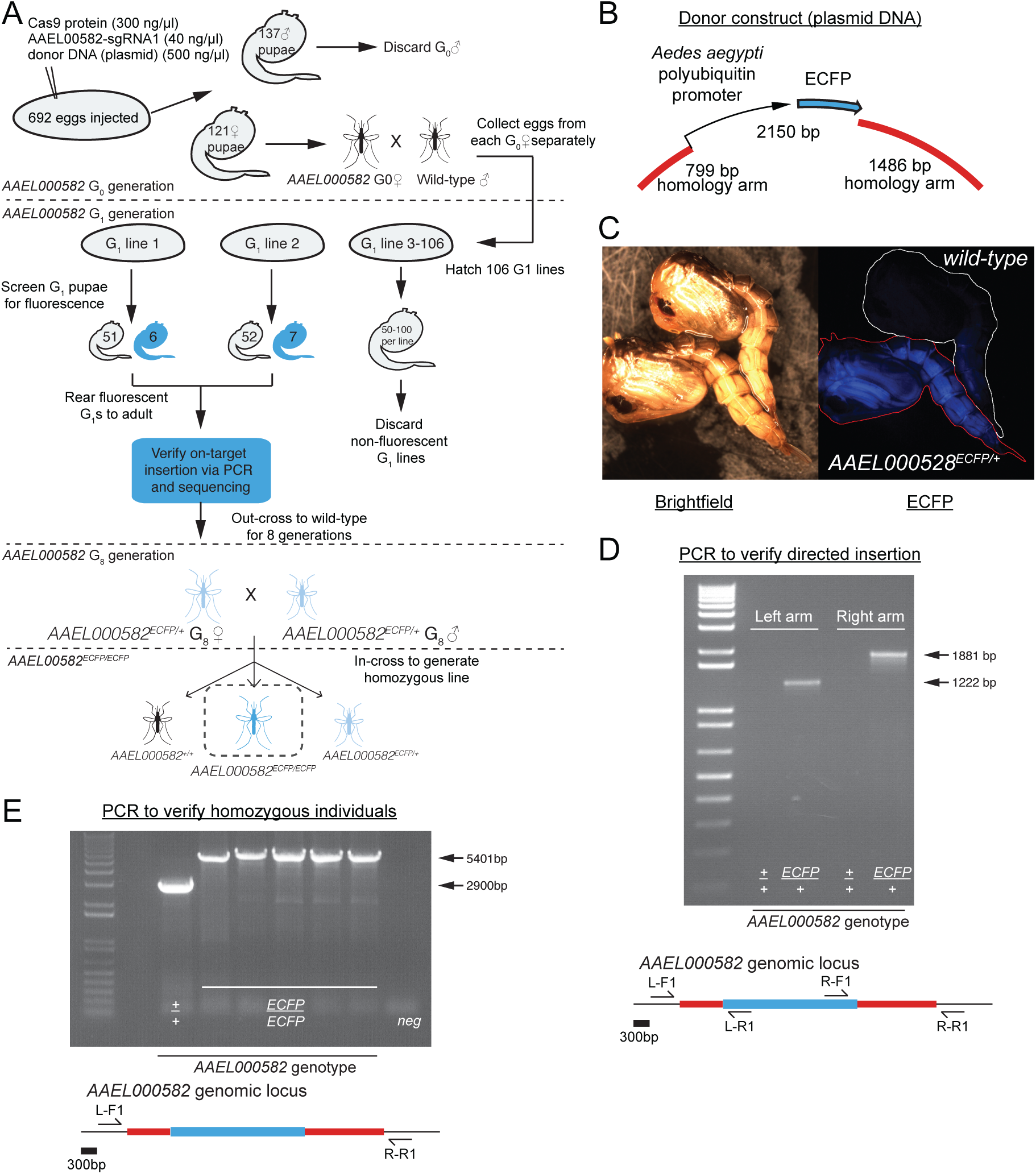
**Insertion of fluorescent cassettes by homology-directed repair A)** Schematic of injection and screening strategy to obtain alleles with an insertion of a fluorescent cassette. Blue pupae represent animals with ECFP expression. **B)** Design of the plasmid donor with homology arms of 799 and 1486 bp surrounding a 2150 bp payload of the PUb promoter driving ubiquitous expression of EFCP. **C)** Brightfield and ECFP fluorescence images of two pupae: wild-type (top) and AAEL000582ECFP/+ (bottom). PUb-ECFP is clearly visible in both larval and pupal stages. **D)** PCR strategy to verify directed insertion of the PUb-ECFP cassette. **E)** PCR strategy to identify homozygous individuals.

G_1_ individuals were screened under a fluorescence dissecting microscope as larvae at 3-5 days post-hatching. The expression of the PUb promoter is clearly visible in larvae and pupae (Figure 6C and 7B). Fluorescent individuals were collected and reared to adulthood and crossed again to wild-type animals to establish stable lines. To verify directed insertion of our cassette as opposed to non-targeted integration at another genomic locus, we designed PCR primers spanning both homology arms (Figure 6D). It is critical that these primers are designed outside each arm and that bands obtained are sequenced to verify junctions between genomic and exogenous sequence on each end of the insertion.

**Figure 7.**
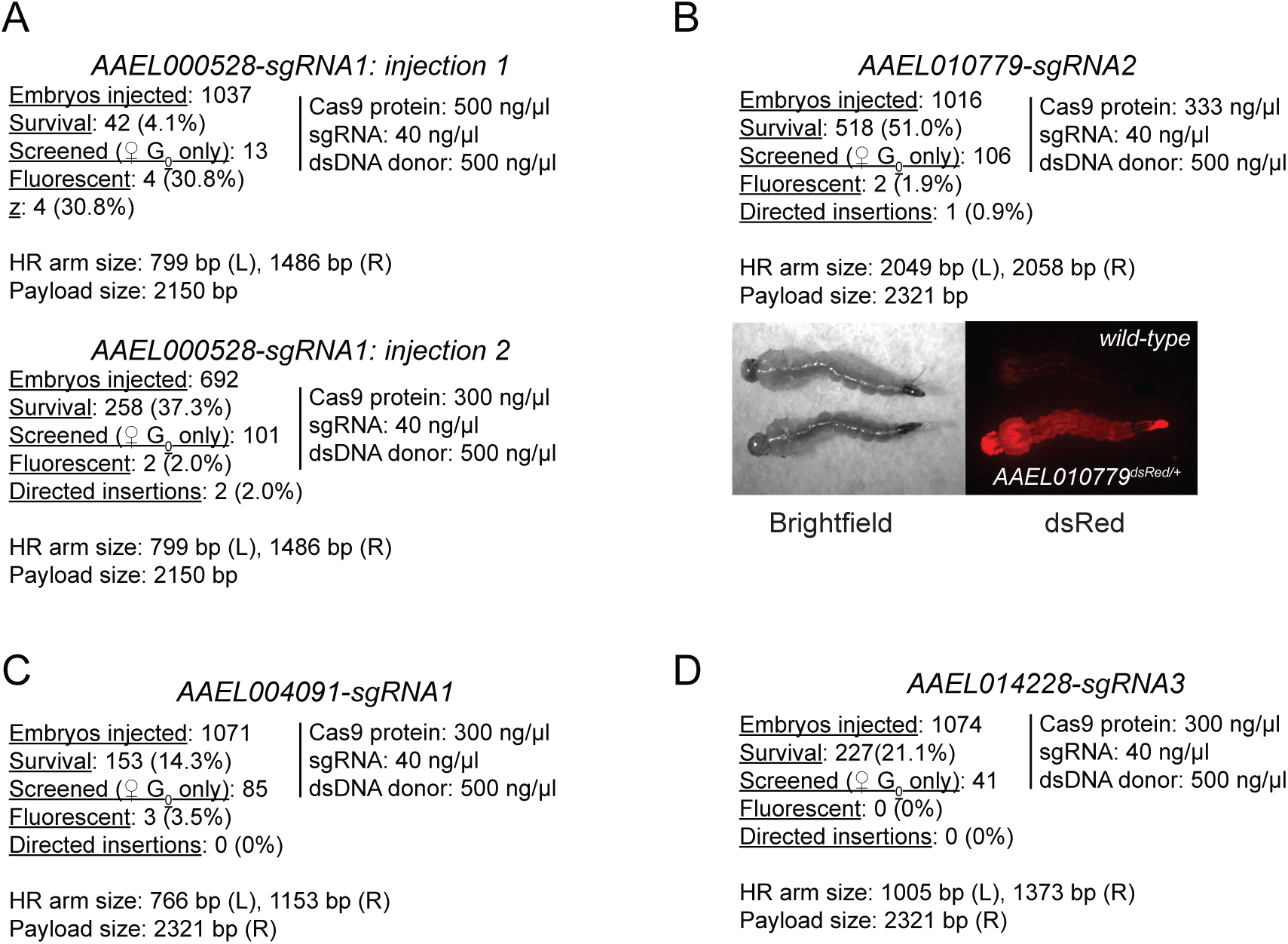
**Summary of injections to insert fluorescent cassettes by homology-directed repair** Table of injection statistics for: **A)** Two successful injections to generate fluorescent insertions in AAEL000582. **B)** One successful injection to generate a fluorescent insertion in AAEL010779. At right, brightfield and dsRed images of a wild-type (top) and AAEL010779dsRed larva (bottom). **C)** One unsuccessful injection attempt to generate a fluorescent insertion in AAEL004091. **D)** One unsuccessful injection attempt to generate a fluorescent insertion in AAEL014228.

Lines containing verified targeted insertions were out-crossed to wild-type mosquitoes for 8 generations, at which point a homozygous line was established by mating heterozygous males and females and selecting putative homozygous individuals from their offspring. Because homozygous individuals show increased levels of fluorescence relative to heterozygotes, the generation of homozygous lines is straightforward. Putative homozygous mosquitoes were separated by sex and used to establish single-pair matings, and the genotype of these single-pair matings were verified by PCR across the integration site of the cassette. In this reaction, PCR primers span the entire cassette, resulting in a large shift in size in homozygous mutant individuals (Figure 6E).

We generated verified targeted insertions in two genomic loci with the *Ae. aegypti* PUb promoter driving the expression of ECFP (Figures 6, 7A) or dsRed (Figure 7B). Attempts at two additional loci failed to produce directed insertion of the donor cassette (Figures 7C-D). We conclude that, similar to the rates of ssODN integration, homology-directed repair from with large plasmid donors occurs at a relatively low frequency compared to other forms of CRISPR-Cas9-mediated genome modification. A drastic variance in the efficiency noted in two injections (Figure 7A) suggests that higher rates can be achieved through simple modifications of injection mix component concentration, perhaps at the expense of embryo survival. The presence of non-directed targeting events underscores the necessity of verifying all lines generated by this technique by PCR or other molecular methods (Figure 6).

Taken together, these data show that the CRISPR-Cas9 system can be been successfully adapted for genome-engineering in the mosquito *Ae. aegypti*. Following the generation of a double-stranded break by an RNA-guided Cas9 nuclease, a variety of mutant alleles can be recovered, including frame-shift mutations caused by insertions or deletions, deletion of a region between two sgRNA target sites, and integration of exogenous sequences from a single-stranded oligonucleotide or a double-stranded plasmid DNA donor. We recommend these methods to any laboratory wishing to generate loss-of-function mutations in *Ae. aegypti*.

## Discussion and Conclusions

We have demonstrated that the CRISPR-Cas9 system is a highly effective and efficient tool for precision genome-editing in the mosquito *Ae. aegypti*. Compared to the relatively low throughput and high cost of ZFN- and TALEN-mediated mutagenesis, the ease and efficiency of designing and producing CRISPR-Cas9 reagents in the laboratory has allowed us to generate stable and precise loss-of-function mutations in over 10 genes to date. This protocol provides a step-by-step manual to mutagenesis in *Ae. aegypti* and also provides general principles that will be useful when translated to other species.

### Optimal injection mix

We recommend using recombinant Cas9 protein for its reproducibility and observed increased rates of mutagenesis and embryo survival. Recombinant Cas9 protein is likely to be much more stable than mRNA, both at the bench and in injected embryos. Additionally, it is likely that sgRNA and Cas9 protein can form a complex in the absence of any other factors, allowing them to generate pre-formed RGENs prior to injection. This can serve to both stabilize sgRNA/Cas9 complexes [33] and ensure that mutagenesis can occur immediately, rather than waiting for Cas9 mRNA to be translated in the embryo. The specific concentrations suggested here represent a good trade-off between survival and efficiency in our hands. However, it is entirely possible that further modifications to this protocol could result in significant increases in certain types of repair.

For CRISPR-Cas9 injections into *Ae. aegypti*, we currently recommend the following injection mix. This mix may also be a good starting point for microinjections into other insect embryos.

- 300 ng/μL recombinant Cas9 protein
- 40 ng/μL sgRNA (each)
- (optional) donor DNA

- 200 ng/μL single stranded ssODN **or**
- 500 ng/μL double stranded plasmid DNA

### Designing active sgRNAs

We observed significant variability in the effectiveness of different sgRNAs, even between sgRNAs targeted to a small genomic region (Figure 3). As in other organisms and cell lines [30, 34], we observed success with sgRNAs ranging in length from 17-20 bp, and recommend considering sgRNAs in this size range when screening for the most active target sites. A single genomic target (AAEL001123) has proven resistant to mutagenesis with 6 different sgRNAs, perhaps reflecting an underlying chromatin-state [35]. Aside from this one example, we were able to design effective sgRNAs to all other examined genes. Systematic examination of sgRNA efficiency in *D. melanogaster* suggests that the GC content of the 6 nucleotides adjacent to the PAM contributes to the efficiency of a given sgRNA [34]. We will incorporate these suggestions as well as those generated from further investigations into the sequence or context-dependency of sgRNA activity into our future sgRNA design. However, given the ease of testing sgRNA efficiency *in vivo*, we still recommend the design and testing of multiple sgRNAs targeting a given gene before committing to large-scale injections.

### Off-target effects

Off-target effects are a concern with any genome-editing technology. We currently adopt several approaches to try and address these concerns in our experiments. First, we check for the sgRNA specificity using publicly available bioinformatic tools [20,36,37], selecting the most specific sgRNAs within the region we wish to target. Second, we design several sgRNAs for a given genomic target. For genes of particular interest, we can generate mutant alleles with multiple sgRNAs and examine their phenotype in transallelic combination, reducing the likelihood that these independent lines will share the same off-target mutations. Third, we have successfully used truncated sgRNAs of less than 20 bp in length, which have been shown in cell culture to reduce the likelihood of off-target modifications [30]. Finally, we subject all lines to at least 8 generations of out-crossing to wild-type mosquitoes, which should reduce the co-inheritance of all but the most tightly linked off-target mutations. While we believe these guidelines reduce the likelihood of significant levels of off-target mutagenesis, we recognize the need for continued efforts to improve and to verify the specificity of all precision genome-engineering technologies.

Variants on the traditional CRISPR-Cas9 reagents designed to improve specificity include the use of paired Cas9 ’nickases’ to generate adjacent single-stranded DNA breaks [38–41]. These dual ’nicks’ are proposed to be less mutagenic than a single double-stranded break. This approach should decrease the chance that the binding of a single sgRNA at an off-target site produces a mutation. Another approach fuses a nuclease-dead Cas9 mutant (dCas9) to the FokI nuclease. As in ZFN and TALENs, FokI requires paired binding, within a certain distance, on opposite strands of a target, and this technique aims to combine the specificity of TALENs and ZFNs with the ease-of-targeting of CRISPR-Cas9 [42,43]. Though we have not investigated these CRISPR-Cas9 variants in our experiments, we believe that their benefits are likely to translate to *Ae. aegypti* and other organisms in the future.

### Enhancing the efficiency of homology-directed repair

In our experiments, insertions and deletions mediated by non-homologous end-joining occur at much higher frequency than by homology-directed repair. This is similar to what has been observed in *Ae. aegypti* with other genome-editing tools, such as ZFNs [2, 11], and in other organisms such as *D. melanogaster* [44].

Several approaches have been developed to increase rates of homology-directed repair. These include injections in the background of a DNA ligase 4 mutation [45,46] or schemes that linearize a double-stranded donor template *in vivo*. Finally, many laboratories working with *D. melanogaster* have developed transgenic strains that express Cas9 protein under ubiquitous or germ-line promoters, improving the efficiency of mutagenesis generally and homology-directed repair specifically [34, 44]. Other areas of interest include alternatives to homology-directed repair, such as ObLiGaRe [47] and homology-independent double-stranded break repair pathways [48]. It remains to be seen whether transgenic Cas9 delivery or alternative integration approaches can be effectively implemented in *Ae. aegypti*.

We note that we have observed a single round of injection that resulted in high (>30%) rates of homology-directed repair but extremely low survival (Figure 7A). This suggests that we might achieve improvements in insertion efficiency by continuing to examine the effects of modulating the concentrations of the three components of the injection mix. If the rates of homology-directed-repair can be sufficiently improved, CRISPR-Cas9 coupled with transgene insertion via homology-directed repair will likely prove to be a versatile tool to tag gene products and introduce transgenes into specific genomic loci, enabling studies of specific neural circuits and other subsets of cells.

### Conclusions

Precision genome-engineering in mosquitoes holds great promise for studies on the genetic basis of behavior [2,11,12] and for genetic strategies to control vector population or disease competence [49]. Ongoing efforts to increase the specificity and efficiency of these technologies is critical to their adaptation as routine techniques, and we believe that the protocols outlined here have met those criteria for the generation of loss-of-function mutations in the mosquito *Ae. aegypti*. Reagents based on the CRISPR-Cas9 platform have been used successfully in organisms from bacteria to primates. This suggests that the techniques described here can likely be adapted to many other non-model organisms, so long as efficient methods of introducing the reagents into the germline and screening for mutations can be developed.

## Acknowledgments

We thank Rob Harrell and the Insect Transformation Facility (University of Maryland) for expert mosquito embryo microinjection, Gloria Gordon and Libby Meija for mosquito rearing assistance, and members of the Vosshall laboratory for helpful comments and critiques. This work was supported in part by contract HHSN272200900039C from the National Institute of Allergy and Infectious Diseases and grant UL1 TR000043 from the National Center for Advancing Translational Sciences (NCATS, National Institutes of Health (NIH) Clinical and Translational Science Award (CTSA) program). B.J.M. was a Jane Coffin Childs Postdoctoral Fellow and L.B.V is an investigator of the Howard Hughes Medical Institute.

